# α2δ-2 is Required for Depolarization-induced Suppression of Excitation in Purkinje cells

**DOI:** 10.1101/2021.03.25.437080

**Authors:** Kathleen A. Beeson, Gary L. Westbrook, Eric Schnell

## Abstract

α2δ proteins (*CACNA2D1-4*) are required for normal neurological function, although how they control neuronal output remains unclear. Using whole-cell recordings of mouse Purkinje cells, we show α2δ-2 is required for functional coupling of postsynaptic voltage-dependent calcium entry with effector mechanisms controlling two different outputs, depolarization-induced suppression of excitation mediated by endocannabinoid signaling, and spike afterhyperpolarization generated by calcium-dependent potassium channels. Our findings indicate an important role for α2δ-2 proteins in regulating functional postsynaptic calcium channel-coupling in neurons.

**Significance Statement:** Calcium influx via membrane voltage-dependent calcium channels drives numerous neuronal processes by signaling through calcium-dependent effector molecules. Signal precision is achieved in part by calcium channel-effector coupling. In mouse Purkinje cell neurons, we show that neuronal α2δ-2 protein functionally couples calcium entry to two different postsynaptic calcium-dependent signals, retrograde endocannabinoid signaling and the action potential afterhyperpolarization. Our findings provide new insights about the control of calcium channel-effector coupling as well as new roles for α2δ-2 proteins in neurons.

## Introduction

Many intracellular signaling cascades are triggered by a common messenger: calcium entering via voltage-gated Ca^2+^ channels (VGCCs) on the plasma membrane. Specificity of Ca^2+^-dependent signaling depends on proximity of VGCCs to effectors in functional nanodomains, and this coupling is critical for neuronal function. For example, in Purkinje cells (PCs), depolarization-induced suppression of excitation (DSE) is initiated via Ca^2+^-dependent endocannabinoid release (1, 2). Likewise, VGCC-K_Ca_ coupling generates spike afterhyperpolarization (AHP), ultimately setting firing frequency (3, 4). Thus, molecules coupling VGCCs to effector-specific signaling critically contribute to transduction of neuronal outputs.

Auxiliary VGCC α2δ proteins (*CACNA2D1-4*) contribute to VGCC membrane trafficking (5), and may couple presynaptic VGCCs to vesicle release machinery (6). However, little is known about auxiliary postsynaptic functions of α2δ outside of these contexts, apart from developmental Ca^2+^-independent roles (7, 8). Here we report that in PCs, which exclusively express α2δ-2 (9, 10), loss of α2δ-2 (*CACNA2D2*) disrupts two disparate forms of postsynaptic VGCC signaling, demonstrating previously unappreciated roles for α2δ proteins in VGCC-effector coupling.

## Results

At PC climbing fiber synapses, postsynaptic VGCC-mediated Ca^2+^ entry initiates retrograde endocannabinoid signaling, acutely reducing presynaptic release probability - a form of plasticity known as DSE (1, 2). Specificity of DSE signaling is achieved by tight functional coupling of postsynaptic VGCCs with Ca^2+^-sensitive endocannabinoid effector molecules (2). Using whole-cell recordings of PCs in acutely prepared brain slices from *CACNA2D2* KO and WT littermate mice, we investigated whether absence of α2δ-2 affects DSE.

In WT PCs, DSE elicited with a 250 ms depolarizing step reduced the amplitude of regularly evoked climbing fiber excitatory postsynaptic currents (EPSCs) by 25% (**Fig 1A-E**). In contrast, DSE was absent in KO PCs (**Fig 1B-E**). The *ducky* mouse, which also lacks α2δ-2 protein, has a reported ∼30% decrease in PC somatic VGCC current density, which is thought to represent decreased VGCC surface trafficking (11). As DSE magnitude is related to the amount of Ca^2+^ influx, it is possible that the reduced VGCC density prevented DSE. However, although increasing depolarizing step length enhances Ca^2+^ influx and DSE in WT mice (1), a four-fold increase in step duration still failed to evoke DSE in α2δ-2 KO PCs **(Fig 1D-F**).

**Figure 1.**
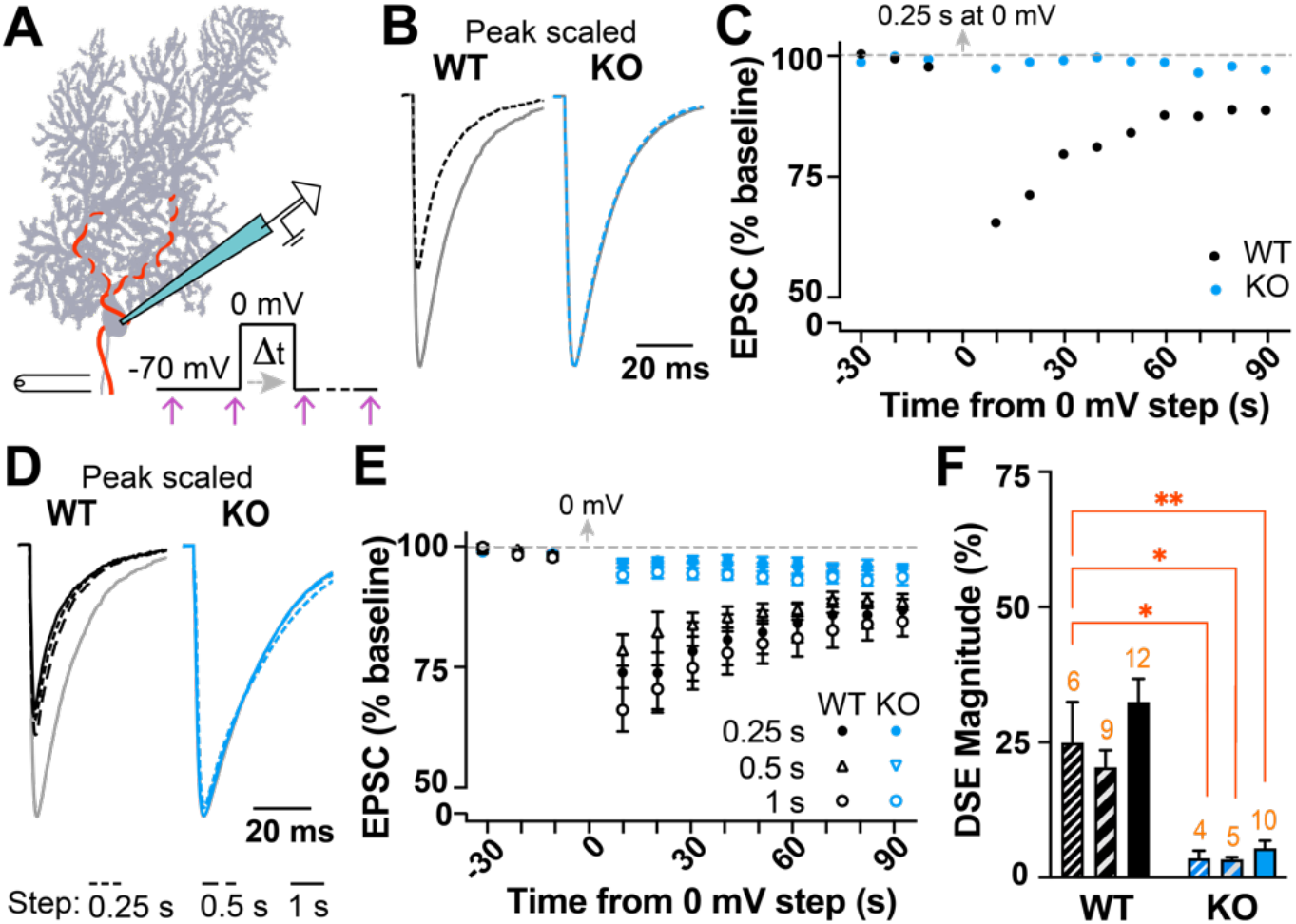
Depolarization-Induced Suppression of Excitation (DSE) reduces climbing fiber-EPSC amplitude in WT but not α2δ-2 KO Purkinje cells. **A)** Experimental schematic. Purkinje cell (PC) is held at −70 mV while the climbing fiber axon (red) is stimulated at 0.2 Hz (arrows). A depolarization step to 0 mV is delivered to the PC between EPSC recordings. **B)** Overlay of peak-scaled EPSC traces during baseline (grey) and 5 s post-depolarization (dotted line) from WT (black) and KO (blue) PCs; 250 ms step duration. **C)** EPSC amplitudes from experiment in B. Each point averaged two consecutive EPSCs. **D)** Overlay of peak-scaled EPSCs during baseline (grey) and 5 s after 250 ms (dotted line), 500 ms (dashed line), and 1 s (solid line) depolarization from WT (black) and KO (blue) PCs. **E)** DSE timecourse from WT (black) and KO (blue) experiments using 250 ms (filled circle), 500 ms (triangle) and 1 s (open circle) depolarizing step lengths. **F)** EPSC depression normalized to baseline in WT (black) and KO (blue) after 250 ms (fine stripe), 500 ms (wide stripe) and 1 s (solid) depolarization steps. n = cells (orange number over bars); each experiment from > 3 mice. One-way ANOVA comparison to average WT 250 ms response, Dunnett’s correction for multiple comparisons; * p < 0.05, ** p < 0.01, *** p < 0.001.

As DSE was not rescued by increased activation of VGCCs in KO PCs, we hypothesized that functional coupling of Ca^2+^ influx to effector molecules was disrupted in the absence of α2δ-2. To examine this, we lowered [EGTA] in our internal solution. As expected for WT PCs, decreasing [EGTA] from 10 mM to either 2 mM or 0.2 mM resulted in more profound DSE magnitude, which further increased with longer voltage steps (**Fig 2A-C**), consistent with greater diffusion of Ca^2+^ from its point of entry. Notably, reduced Ca^2+^ buffering restored DSE in KO PCs (**Fig 2D-F**), indicating that signaling mechanisms involved in DSE expression remained intact. Thus, these data indicate that rather than affecting Ca^2+^ entry per se, α2δ-2 mediates tight functional coupling between VGCCs and endocannabinoid release.

**Figure 2.**
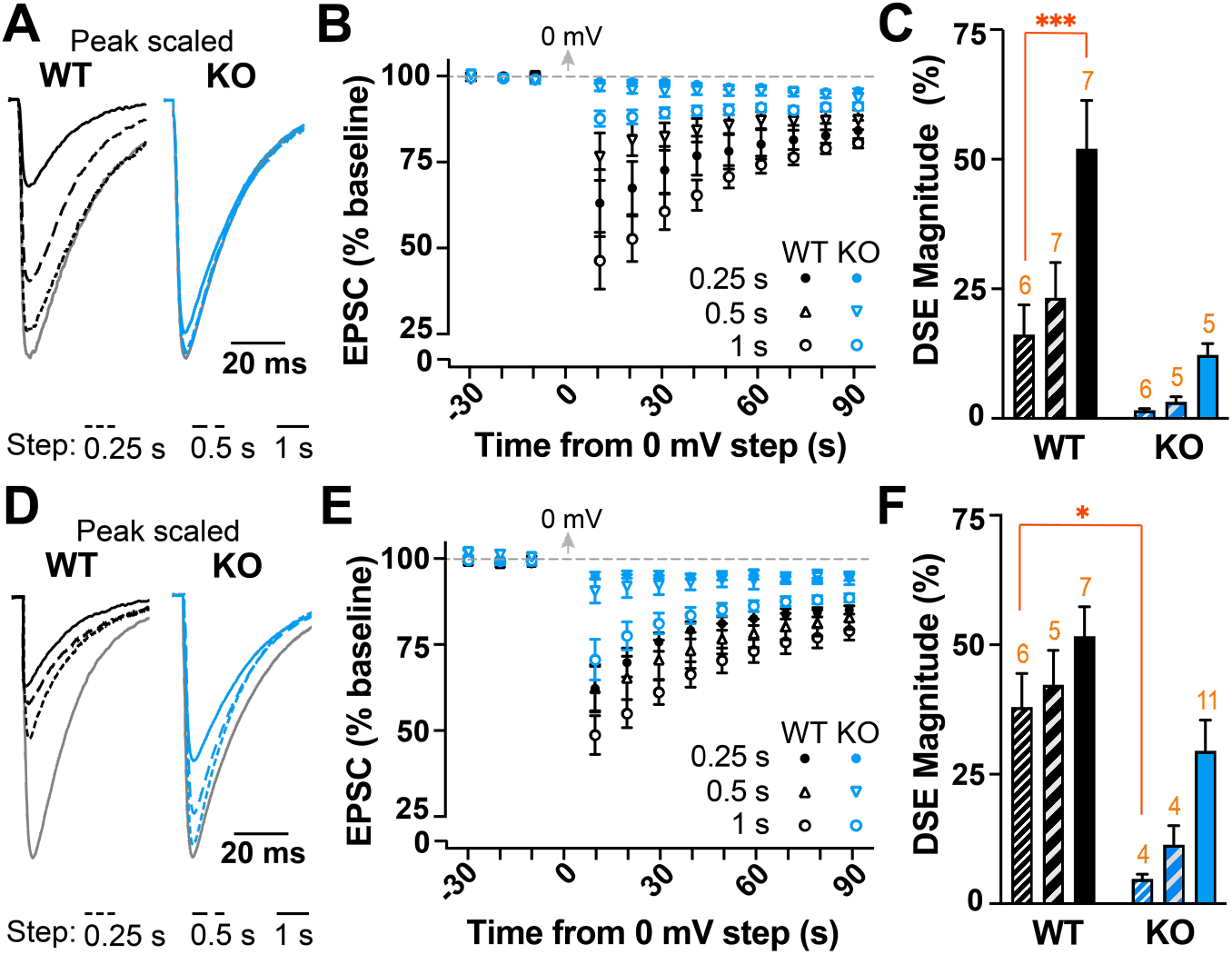
Reduced intracellular [EGTA] reveals DSE in the α2δ-2 KO. **A, D**) Overlay of peak-scaled EPSCs during baseline (grey) and 5 s after 250 ms (dotted line), 500 ms (dashed line), and 1 s (solid line) depolarization using intracellular solution containing 2 mM (**A**) or 0.2 mM (**D**) EGTA from WT (black) and KO (blue) PCs. **B, E**) DSE timecourse from WT (black) and KO (blue) experiments using a 2 mM (**B**) or 0.2 mM (**E**) EGTA and depolarization steps of 250 ms (filled circle), 500 ms (triangle) and 1 s (open circle). **C, F**) EPSC depression normalized to baseline in WT (black) and KO (blue) after 250 ms (fine stripe), 500 ms (wide stripe) and 1 s (solid) depolarization steps using a 2 mM (**C**) or 0.2 mM (**F**) EGTA. n = cells (orange number over bars); each experiment from > 3 mice. One-way ANOVA comparison to average WT 250 ms response, Dunnett’s correction for multiple comparisons; * p < 0.05, ** p < 0.01, *** p < 0.001.

To resolve whether α2δ-2 affects coupling of other molecules to postsynaptic VGCC nanodomains, we focused on Ca^2+^-dependent action potential AHPs. In PCs, the AHP is mediated by BK-type K_Ca_ channels (12, 13), and regulates PC firing rate (3, 4). To assess VGCC-K_Ca_ coupling, we recorded spontaneous spiking in PCs from WT and KO mice during whole-cell current-clamp recordings. In agreement with previous studies in *ducky* mutants (11, 14), tonic spike rate in α2δ-2 KO PCs was reduced compared to WT (**Fig 3A-B**). Furthermore, AHP amplitude of individual spike waveforms was consistently smaller in α2δ-2 KO cells (**Fig 3C-D**), indicating reduced K_Ca_ channel activation (4, 12). Other membrane properties were unchanged in KO PCs, including the resting membrane potential, membrane polarization rate, spike threshold and spike height (**Fig 3E-J**), indicating that changes were limited to the K_Ca_-mediated AHP. Additionally, there was no change in BK membrane localization in PCs by immunohistochemistry (**Fig 3K**; Membrane BK density (puncta/μm): WT = 0.82 ± 0.13, n = 4; KO = 0.75 ± 0.08, n = 3; p = 0.7; Student’s unpaired t-test), consistent with normal BK expression and function, as previously described in *ducky* mice (15).

**Figure 3.**
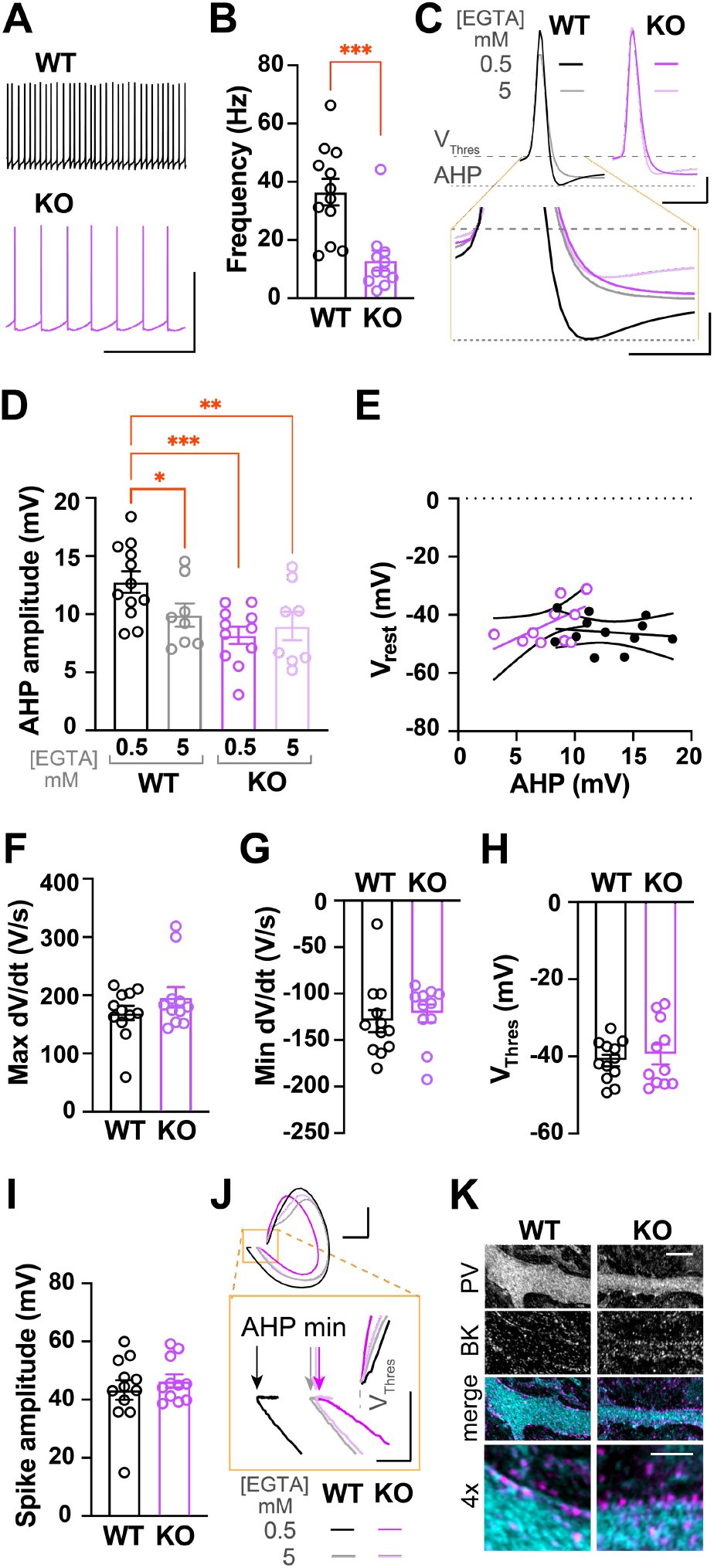
Spontaneous firing frequency, afterhyperpolarization (AHP) amplitude and Ca^2+^ coupling are reduced in the α2δ-2 KO. **A)** Spontaneous spikes in WT (black) and KO (magenta) PCs; scale: 50 mV, 0.5 s. **B)** Spontaneous firing frequency in PCs. Student’s unpaired t-test, p < 0.0001. **C)** Averaged spontaneous spikes from WT and KO PCs during tonic firing (0.5 mM EGTA, black/magenta; 5 mM EGTA grey/light purple); scale: 10 mV, 2 ms. Grey dotted lines indicate V_thres_ and minimum voltage during AHP. *Below*, enlarged overlay demonstrating differences in AHP amplitude; scale: 5 mV, 1 ms. **D)** AHP amplitudes recorded using 0.5 mM EGTA or 5 mM EGTA intracellular solution. One-way ANOVA comparison to average WT 0.5 mM EGTA response, uncorrected Fischer’s LSD; * p < 0.05, ** p < 0.01, *** p < 0.001. **E)** Correlation of AHP amplitude vs. resting membrane potential (V_m_) in WT (black) and KO (magenta) PCs using 0.5 mM EGTA. Linear regression and 95% confidence interval; WT R^2^ = <0.002; KO R^2^ = 0.393. **F-I**) Spike waveform parameters in WT and KO PCs. No differences in maximum dV/dt (**F**), minimum dV/dt (**G**), spike threshold (V_thres_) (**H**) or spike height (**I**); unpaired Student’s t-tests. **J)** Phase plane plots of spontaneous spikes in WT and KO PCs (0.5 mM EGTA, black/magenta; 5 mM EGTA grey/light purple). Traces aligned by spike threshold (V_thresh_) for comparison; scale: 100 mV/ms, 20 mV. *Below*, enlarged inset to illustrate differences in AHP minimum amplitude (arrows; 0.5 mM EGTA, black/magenta; 5 mM EGTA, grey/light purple); scale: 100 mV/ms, 5 mV. **K)** Immunohistochemistry of WT and KO cerebellar slices stained for parvalbumin (PV; cyan) and the BK channel (magenta); scale: 5 µm. *Below*, merged higher power image; scale: 2 µm.

To determine whether the reduced AHP resulted from functionally uncoupled VGCC-K_Ca_ channels in KO mice, we dialyzed PCs with an increased EGTA concentration (from 0.5 mM to 5 mM) to uncouple VGCC-BK signaling (16). This additional Ca^2+^ buffering reduced the AHP amplitude in WT PCs to the same value as the KO, with no effect on KO AHP (**Fig 3C-D**). Thus, increased Ca^2+^ buffering uncoupled VGCC-K_Ca_ signaling in WT, but K_Ca_ channels were already functionally uncoupled in the α2δ-2 KO.

## Discussion

As a primary signal in neurons, precise spatiotemporal regulation of Ca^2+^ influx maintains fidelity of Ca^2+^-dependent processes in neurons. Consequently, molecules controlling VGCC coupling to downstream effectors are critical to neuronal function. α2δ proteins are involved trafficking of VGCCs to the plasma membrane (5). However, this conclusion was necessarily based on heterologous expression systems, as most neurons express more than one α2δ isoform. Thus, information of postsynaptic auxiliary functional roles of α2δ proteins in neurons is limited. Because PCs selectively express the α2δ-2 isoform (9, 10), the *CACNA2D2* KO mouse provides an ideal model to examine the functional roles for α2δ proteins in the postsynaptic compartment. Our results demonstrate that two distinct Ca^2+^-dependent mechanisms, DSE and K_Ca_ signaling, are disrupted in α2δ-2 KO PCs, indicating functional loss of postsynaptic VGCC-effector coupling (2, 16).

How does α2δ-2, a largely extracellular protein, mediate functional coupling of VGCCs with intracellular effector proteins? It is possible that α2δ-2 directly associates with other extracellular proteins involved in VGCC domains (5). As endocannabinoid machinery resides at synapses (17), and α2δ proteins are important for synapse formation (7, 9, 10, 18), α2δ could potentially regulate VGCC nanodomains by binding to presynaptic adhesion proteins. Another possibility is that α2δ-2 localizes VGCCs to lipid rafts. Though VGCCs are abundant in non-lipid raft membrane fractions where they are independent of α2δ-2, VGCCs and α2δ-2 colocalize in lipid rafts isolated from cerebellar homogenates (19). Intriguingly, DAG lipase-α, the enzyme responsible for synthesis of endocannabinoids involved in DSE, has also been isolated in lipid rafts (17), and mislocalization of VGCCs away from lipid rafts might explain the reduced efficacy of VGCC-signaling in α2δ-2 KO PCs. The deficits in Ca^2+^-dependent signaling we observed may contribute to the dramatic neurologic phenotypes in α2δ-2 KO mice. Our results provide clues for future work to directly assay α2δ-2 interacting proteins and subcellular localization of VGCCs in PCs. Future investigations of the coupling roles of other α2δ isoforms in neurons will provide valuable insights into how these proteins impact neurological functions across the brain.

## Materials and Methods

### Animals

*Cacna2d2* knockout mice (*Cacna2d2*^*tm1Svi*^, MGI = 3055290; generously supplied by Drs. Sergey Ivanov and Lino Tessarollo) were obtained as cryopreserved sperm and re-derived via *in vitro* fertilization on a C57BL/6J background. Breeding mice were kept heterozygous, and genotyping was performed as previously described (10). Mice were maintained in facilities fully accredited by the Association for Assessment and Accreditation of Laboratory Animal Care and veterinary care was provided by Oregon Health & Science University’s Department of Comparative Medicine. All animal care and experiments were performed in accordance with state and federal guidelines, and all protocols were approved by the OHSU Institutional Animal Care and Use Committee.

### Slice Preparation and Electrophysiology

Male and female mice were used between the ages of p21-30. KO and WT littermates were deeply anesthetized and transcardially perfused with ice-cold choline-based solution containing (mM): 125 choline-Cl, 2.5 KCl, 1.25 NaH_2_PO_4_, 0.44 ascorbate, 2 Na pyruvate, 3 3-myo-inositol, 10 D-glucose, 25 NaHCO_3_, 7 MgCl_2_, 0.5 CaCl_2_ (osmolarity adjusted to 305 mOsm) and equilibrated with 95% O_2_ and 5% CO_2_ gas mixture. Acute 300 µm sagittal slices were cut from cerebellum using a vibratome (VT1200, Leica Microsystems), and incubated for 30 minutes in standard artificial cerebral spinal fluid (aCSF) at 34°C.

### Voltage clamp recordings

Whole-cell recordings were obtained using 1-3 MΩ borosilicate glass pipettes filled with internal solution containing (in mM): 100 CsMeSO_4_, 35 CsCl, 15 TEA-Cl, 1 MgCl_2_, 15 HEPES, 2 ATP-Mg, 0.3 TrisGTP, 10 phosphocreatine, and 2 QX-314. A large batch of this internal base solution was equally divided and 10, 2 or 0.2 mM EGTA was added to each third. All internals were adjusted to pH 7.3 with CsOH and osmolarity to 293 mOsm. External solution contained (in mM): 125 NaCl, 25 NaHCO_3_, 1.25 NaH_2_PO_4_, 3 KCl, 25 Dextrose, 2 CaCl_2_, 1 MgCl_2_ (osmolarity adjusted to 300 mOsm) and was continuously perfused via roller pump.

PCs were identified and recorded as previously described (10). Briefly, PCs were chosen from the vermis lobe VI, were identified by soma size and location in the PC layer, and whole-cell patch-clamp recordings were obtained in voltage clamp mode. Cell capacitance, series resistance and input resistance monitored in real time using intermittent −10 mV voltage steps. Inhibition was blocked in all experiments by 10 µM SR95531 (Tocris), and 0.2-0.5 µM NBQX (Tocris) was included to maintain voltage clamp of climbing fiber-mediated excitatory postsynaptic currents (EPSCs). All voltage clamp recordings were performed at room temperature. Signals were amplified with a MultiClamp 700B (Molecular Devices) amplifier and pipette capacitance was compensated using MultiClamp software. Signals were low-pass filtered at 6 kHz and sampled at 10 kHz, and digitized with a National Instruments analog-to-digital board. All recordings were acquired and analyzed using IgorPro-based (Wavemetrics) software.

For DSE experiments, PCs were held at −70 mV while climbing fiber-mediated EPSCs were evoked using a monopolar glass electrode in the granule cell layer. After obtaining 2 minutes of baseline responses at 0.2 Hz, a depolarizing voltage step to 0 mV of 1 s, 500 ms or 250 ms duration was delivered to induce DSE, after which PCs were returned to −70 mV and 0.2 Hz stimulation was continued. DSE plasticity is acute, and most synapses recover back to baseline EPSC amplitudes within < 60 seconds (1). As a small amount of “run down” was routinely observed in the evoked CF-mediated EPSC amplitude, DSE inclusion criteria required EPSC amplitudes to return to 80% baseline within 2 minutes post-stimulation (opposed to “stepping” to decreased amplitude without recovery). A minimum of 5 minutes were waited between DSE inductions, and step length was randomized throughout experiment. Series resistance was not compensated; cells with series resistance >10 MΩ, or a >2 MΩ change in series resistance over the course of the experiment were excluded.

For analysis, EPSC amplitudes were binned every 10 seconds (2 traces) and normalized to the 1 minute of baseline immediately preceding the depolarizing step. The ‘DSE magnitude’ (e.g. Figure 2E) is based on the average of EPSC amplitude 5 and 10 s after the depolarizing step. Example traces shown are from 5 s after the depolarizing step. A minimum of 3 mice per genotype were used for each manipulation, with no more than 2 cells/treatment coming from one mouse. For data presentation, EPSC traces were off-line box-filtered at 1 kHz in Igor64 software.

### Current clamp recordings

For spontaneous spike experiments, internal solution contained (in mM): 120 KCH_3_SO_3_, 10 HEPES, 10 NaCl, 2 MgCl_2_, 0.5 EGTA, 4 ATP-Mg, 0.3 Tris-GTP, and 14 phosphocreatine, pH 7.35 adjusted with KOH (osmolarity adjusted to 293 mOsm). A stock of EGTA solution was added to aliquots of internal, to increase [EGTA] to 5mM as needed. Synaptic inhibition was achieved with 10 µM SR95531 (Tocris) and 10 µM NBQX (Tocris), and recordings were made at 36°C using an in-line heater. PCs in whole-cell mode from vermis lobe VI were first held in voltage clamp mode to monitor access series and input resistance before switching to current clamp. Changes in access were corrected with bridge balance using Multiclamp software. For increased action potential waveform resolution, some current clamp experiments were sampled at 50 kHz.

Spontaneous spikes from tonically firing PCs with < 400 pA holding current and < 10 MOhm series were analyzed using the Igor64 Neuromatic tools. Firing frequency data was collected from 10 seconds of recording, which yielded ∼100-500 spikes. Action potential properties were assessed by averaging 50 consecutive spikes. Afterhyperpolarization (AHP) amplitude was measured as the difference between the threshold voltage (V_thres_ = depolarization rate >10 V/s) and the minimum voltage reached within 5 ms of spiking. All current clamp data was taken at least 3 minutes after break-in to allow time for internal solution to dialyze. Spike traces were box filtered for data visualization, and phase plane plots were made using Igor64.

### Immunohistochemistry

Immunohistochemistry was performed as described (10). Briefly, p21 WT and KO mice were deeply anesthetized and transcardially perfused with ice-cold PBS followed by 4% paraformaldehyde (PFA)-PBS. Following decapitation, brains were removed and fixed overnight in 4% PFA-PBS, and stored in PBS at 4°C. Sagittal cerebellar slices were made at 50 µm thickness using a vibratome, and slices containing vermis lobe VI were permeabilized for 1 hr with 0.4% Triton-PBS with 10% normal horse serum at RT. Slices were stained with goat anti-Parvalbumin (Swant #PVG-213; 1:1000) and mouse anti-BK (Neuromab #73-022; 1:500) overnight at 4°C. Corresponding fluorescently labeled secondaries (Invitrogen; 1:500) were applied after rinsing 3x in PBS, and slices were mounted on glass cover slips using Fluoromount G (Sigma-Aldrich).

BK membrane expression was imaged using 63x oil immersion lens on a LSM980 microscope with ZEN software. ∼7 µm z-stack images of primary PC dendrites were acquired at 0.15 µm intervals using the PV channel at 4.5 x zoom with 680 × 680 pixel resolution. Airyscan images were processed using default settings in ZEN. Quantification of membrane localized BK puncta was done by a separate researcher, blinded to genotype, using the most transverse section of dendrite from each z-stack. For presentation, images were processed in Fiji/ImageJ, illustrating 0.45 µm maximum projections, and the panel was assembled using Adobe Photoshop.

### Statistics

The data were tested for normality using Shapiro-Wilk test. Data from male and female mice were grouped. The difference in magnitude of DSE between the WT 250 ms depolarization step condition and other groups were compared using a one-way ANOVA with Dunnett’s correction for multiple comparisons. For current clamp data, student’s unpaired t-tests were used for spike frequency comparison between WT and KO. In current clamp experiments using 5 mM EGTA, only AHP amplitude was significantly different (all other measures not shown). For this data, a one-way ANOVA with Fisher’s LSD was used to compare all groups to WT 0.5 mM condition. All electrophysiology experiments utilized at least 3 animals per genotype, where n = # cells. For immunohistochemistry of BK membrane density measurements, 2-4 images per animal were averaged (n = mice), and an unpaired t-test was used for comparison. Data were graphed in Prism GraphPad version 8 and are reported as the mean ± SEM. *p<0.05, **p<0.01, ***p<0.001.

## Acknowledgments

This research was supported by VA I01-BX004938 (ES), DoD W81XWH-18-1-0598 (ES), NIH T32NS007466 (KAB), NIH R01-NS080979 (GLW), NIH P30NS061800 (OHSU), and NINDS 1R21NS102948 (Ines Koerner/ES). The contents of this manuscript do not represent the views of the U.S. Department of Veterans Affairs or the United States Government.

